# Integration of experiments across diverse environments identifies the genetic determinants of variation in *Sorghum bicolor* seed element composition

**DOI:** 10.1101/019083

**Authors:** Nadia Shakoor, Greg Ziegler, Brian P. Dilkes, Zachary Brenton, Richard Boyles, Erin L. Connolly, Stephen Kresovich, Ivan Baxter

## Abstract

**Abstract:** Seedling establishment and seed nutritional quality require the sequestration of sufficient element nutrients. Identification of genes and alleles that modify element content in the grains of cereals, including *Sorghum bicolor*, is fundamental to developing breeding and selection methods aimed at increasing bioavailable element content and improving crop growth. We have developed a high throughput workflow for the simultaneous measurement of multiple elements in sorghum seeds. We measured seed element levels in the genotyped Sorghum Association Panel (SAP), representing all major cultivated sorghum races from diverse geographic and climatic regions, and mapped alleles contributing to seed element variation across three environments by genome-wide association. We observed significant phenotypic and genetic correlation between several elements across multiple years and diverse environments. The power of combining high-precision measurements with genome wide association was demonstrated by implementing rank transformation and a multilocus mixed model (MLMM) to map alleles controlling 20 element traits, identifying 255 loci affecting the sorghum seed ionome. Sequence similarity to genes characterized in previous studies identified likely causative genes for the accumulation of zinc (Zn) manganese (Mn), nickel (Ni), calcium (Ca) and cadmium (Cd) in sorghum seed. In addition to strong candidates for these four elements, we provide a list of candidate loci for several other elements. Our approach enabled identification of SNPs in strong LD with causative polymorphisms that can be evaluated in targeted selection strategies for plant breeding and improvement.

**One sentence summary:** High-throughput measurements of element accumulation and genome-wide association analysis across multiple environments identified novel alleles controlling seed element accumulation in *Sorghum bicolor.*

This project was partially funded by the iHUB Visiting Scientist Program (http://www.ionomicshub.org), Chromatin, Inc., NSF EAGER (1450341) to I.B. and BPD, NSF IOS 1126950 to IB, NSF IOS-0919739 to EC, and BMGF (OPP 1052924) to B.P.D.

## Introduction

*Sorghum bicolor* is a globally cultivated source of food, feed, sugar and fiber. Classified as a bioenergy feedstock, sorghum biomass also has unique advantages for sustainable biofuel production (Kimber et al., 2013). The element composition of stems, leaves and reproductive organs all contribute significantly to biomass quality. The seed bearing reproductive organs, or panicles, in sorghum represent up to 30% of the total dry matter yield (Amaducci et al., 2004). Inorganic elements, particularly alkali metals, influence the combustion process and can limit the effectiveness of biomass conversion (Obernberger et al., 1997; Monti et al., 2008). Targeted reduction of specific elements and compositional traits via transgenic and breeding approaches can be implemented to improve biomass quality.

Increasing the bioavailable elemental nutrient content in the edible portions of the crop has the potential to increase the value of sorghum for human and animal nutrition. Plant-based diets, in which grains compose the major food source, require the availability of essential elements in the plant seed. Iron (Fe) and Zn deficiencies affect over 2 billion people worldwide (Organization, 2002), and increases in the accumulation and bioavailability of these elements in cereal grains, including sorghum, could potentially make a significant impact towards ameliorating this nutritional crisis (Graham et al., 1999; Organization, 2002). Additional global health benefits could be achieved by increasing magnesium (Mg), selenium (Se), Ca and copper (Cu) (White and Broadley, 2005) and reducing the concentration of toxic elements, including arsenic (As) and Cd (Ma et al., 2008).

Seed element content is determined by interconnected biological processes, including element uptake by the roots, translocation and remobilization within the plant, and ultimately import, deposition and assimilation/storage in the seeds. Element availability is further affected by the accumulation of metabolites in seeds (Vreugdenhil et al., 2004). High-throughput ionomic analysis, or concurrent measurement of multiple elements, allows for the quantitative and simultaneous measurement of an organism’s elemental composition, providing a snapshot into the functional state of an organism under different experimental conditions (Salt et al., 2008). Most studies of the plant ionome utilize inductively-coupled plasma mass spectroscopy (ICP-MS). Briefly, the ICP functions to ionize the analyte into atoms, which are then detected by mass spectroscopy. Reference standards are utilized to quantify each element of interest in the sample analyte. ICP-MS analysis time is approximately 1–3 minutes per sample, which allows for a high-throughput processing of hundreds of samples (Salt et al., 2008). Previous studies have demonstrated that several elements, including Fe, Mn, Zn, cobalt (Co) and Cd share mechanisms of accumulation (Yi and Guerinot, 1996; Vert et al., 2002; Connolly et al., 2003). Ionomics signatures derived from multiple elements have been shown to better predict plant physiological status than the measurements of the elements themselves, including the essential nutrients (Baxter et al., 2008). Holistically examining the ionome provides significant insights into the networks underlying ion homeostasis beyond single element studies (Baxter and Dilkes, 2012).

With over 45,000 catalogued sorghum germplasm lines (USDA), there is significant genetic variation (Das et al., 1997) with undiscovered impact on seed element composition. Mapping quantitative trait loci (QTL) for seed element concentration has been successful in a number of species including Arabidopsis (Vreugdenhil et al., 2004; Waters and Grusak, 2008; Buescher et al., 2010), rice (Norton et al., 2010; Zhang et al., 2014), wheat (Shi et al., 2008; Peleg et al., 2009) and maize (Šimić et al., 2012; Baxter et al., 2013; Baxter et al., 2014). Genome-wide association (GWA) mapping is well suited for uncovering the genetic basis for complex traits, including seed element accumulation. One of the key strengths of association mapping is that *a priori* knowledge is not necessary to identify new loci associated with the trait of interest. Further, a GWA mapping population is comprised of lines that have undergone numerous recombination events, allowing for a narrower mapping interval. Previous GWA studies in maize (Tian et al., 2011), rice (Huang et al., 2010) and sorghum (Morris et al., 2013) have been successful in identifying the genetic basis for various agronomic traits. Here, we analyzed the seed ionome from a community-generated association panel to identify potential loci underlying seed element accumulation in sorghum.

## Results

### Phenotypic diversity for seed element concentrations in the sorghum association panel

We grew 407 lines from the publicly available sorghum association panel (SAP) selected for genotypic diversity and phenotypic variation (Casa et al., 2008) (Supplemental table 1). These lines were previously genotyped by sequencing (GBS) (Morris et al., 2013). The SAP lines were grown in three experiments: Lubbock, Texas in 2008 (SAP 2008), Puerto Vallarta, Mexico in 2012 (SAP 2012), and two field replicates produced in Florence, SC in 2013 (SAP 2013-1 and SAP2013-2). 287 of the 407 SAP lines were present in all 4 growouts.

Seed samples were taken from each replicate and weighed before analysis. A simple weight normalization and established methods to estimate weight from the elemental content were attempted (Lahner et al., 2003). However, both methods created artifacts, particularly in elements with concentrations near the level of detection (Supplemental figure 1). To address this concern, we included weight as a cofactor in a linear model that included other sources of technical error and utilized the residuals of the model as the trait of interest for genetic mapping. The residuals from this transformation were used for all further analyses and outperformed any other method (data not shown).

We calculated broad-sense heritability for each trait to determine the proportion of the phenotypic variation explained by the genetic variation present in the SAP across the three environments (Table 1). Heritability estimates ranged from 1% (sodium, Na) to 45% (Cu). We obtained moderate heritability (> 30%) for several elements including: Mg, P, sulfur (S), potassium (K), Ca, Mn, Fe, Co, Zn, strontium (Sr) and molybdenum (Mo). Low heritabilities were reported previously for seed accumulation of Al and As (Norton et al., 2010) as well as for Se, Na, Al, and Rb in a similarly designed study in maize seed kernels (Baxter et al., 2014). The relatively lower heritabilities for these elements, including boron (B), Cd and Se could be explained by environmental differences between the experiments, element accumulation near the limits of detection via ICP-MS, or the absence of genetic variation affecting these element’s concentrations. Consistent with the hypothesis that field environment was masking genetic variation, we calculated the heritability for two field replicates of the SAP in 2013, and found higher heritabilities for 12 elements (Table 1).

**Table 1.** Mean, standard deviation, and broad sense heritability of seed element concentrations from the Sorghum Association Panel averaged across 3 environments. Element concentration values are presented as mg kg^-1^ and broad sense heritability (H^2^) was calculated as described in the methods section. Data represents an average of individual samples (n=287) analyzed in 4 separate experiments. *Element concentration presented in μg kg^−1^.

| Trait | Sorghum Association Panel |  | H<sup>2</sup> | H<sup>2</sup> |
| --- | --- | --- | --- | --- |
|  | All SAP Replicates |  | All SAP | 2013 Field |
|  | Mean | Standard Deviation | Reps | Reps |
| **B** | 12.8 | 5.88 | 0.02 | 0.05 |
| **Na** | 0.572 | 0.316 | 0.01 | 0.00 |
| **Mg** | 1580 | 246 | 0.42 | 0.38 |
| **Al** | 0.277 | 0.269 | 0.06 | 0.17 |
| **P** | 3350 | 623 | 0.38 | 0.35 |
| **S** | 890 | 127 | 0.36 | 0.33 |
| **K** | 3850 | 795 | 0.36 | 0.46 |
| **Ca** | 25.3 | 18.8 | 0.41 | 0.57 |
| **Mn** | 13.5 | 3.33 | 0.36 | 0.43 |
| **Fe** | 24.9 | 5.7 | 0.40 | 0.46 |
| **Co** | 6.11* | 4.48* | 0.32 | 0.28 |
| **Ni** | 0.18 | 0.187 | 0.22 | 0.10 |
| **Cu** | 2.54 | 1.21 | 0.45 | 0.47 |
| **Zn** | 19.9 | 5.71 | 0.35 | 0.38 |
| **As** | 0.0796 | 0.0361 | 0.08 | 0.19 |
| **Se** | 1.32 | 0.454 | 0.03 | 0.00 |
| **Rb** | 1.76 | 0.848 | 0.10 | 0.12 |
| **Sr** | 0.573 | 0.586 | 0.32 | 0.27 |
| **Mo** | 0.603 | 0.306 | 0.33 | 0.37 |
| **Cd** | 0.0683 | 0.0739 | 0.16 | 0.23 |

We detected significant effects of both genotype and environment on most of the elements (Figure 1 and Supplemental table 2). The measured element concentrations of the present study corroborate the broad range observed in the sorghum element literature (Mengesha, 1966; Neucere and Sumrell, 1980; Lestienne et al., 2005; Ragaee et al., 2006). Similar to a study carried out in wild emmer wheat (Gomez-Becerra et al., 2010), grain Na and Ca showed large variation (5 and 4 fold, respectively). Compared to micronutrients, the remaining macronutrients (P, K, S and Mg) measured in the study exhibited less phenotypic variation overall (Table 1 and Supplemental table 3) ranging between 1.6 and 1.8 fold across the SAP. Of the micronutrients, high variation was detected for Al and Ni (8 and 6 fold, respectively). With the exception of these two elements, seed micronutrient concentration showed phenotypic variation ranging between 2.4 to 5.6 fold. High variation in Ni and Al may indicate strong environmental effects on grain Ni and Al concentration or contamination during handling and analysis of the seeds, as previously suggested (Baxter et al., 2014). The element traits were well distributed across the sorghum subpopulations, with no specific subpopulations accumulating disproportionate levels of any element (Supplemental figure 2).

**Figure 1.**
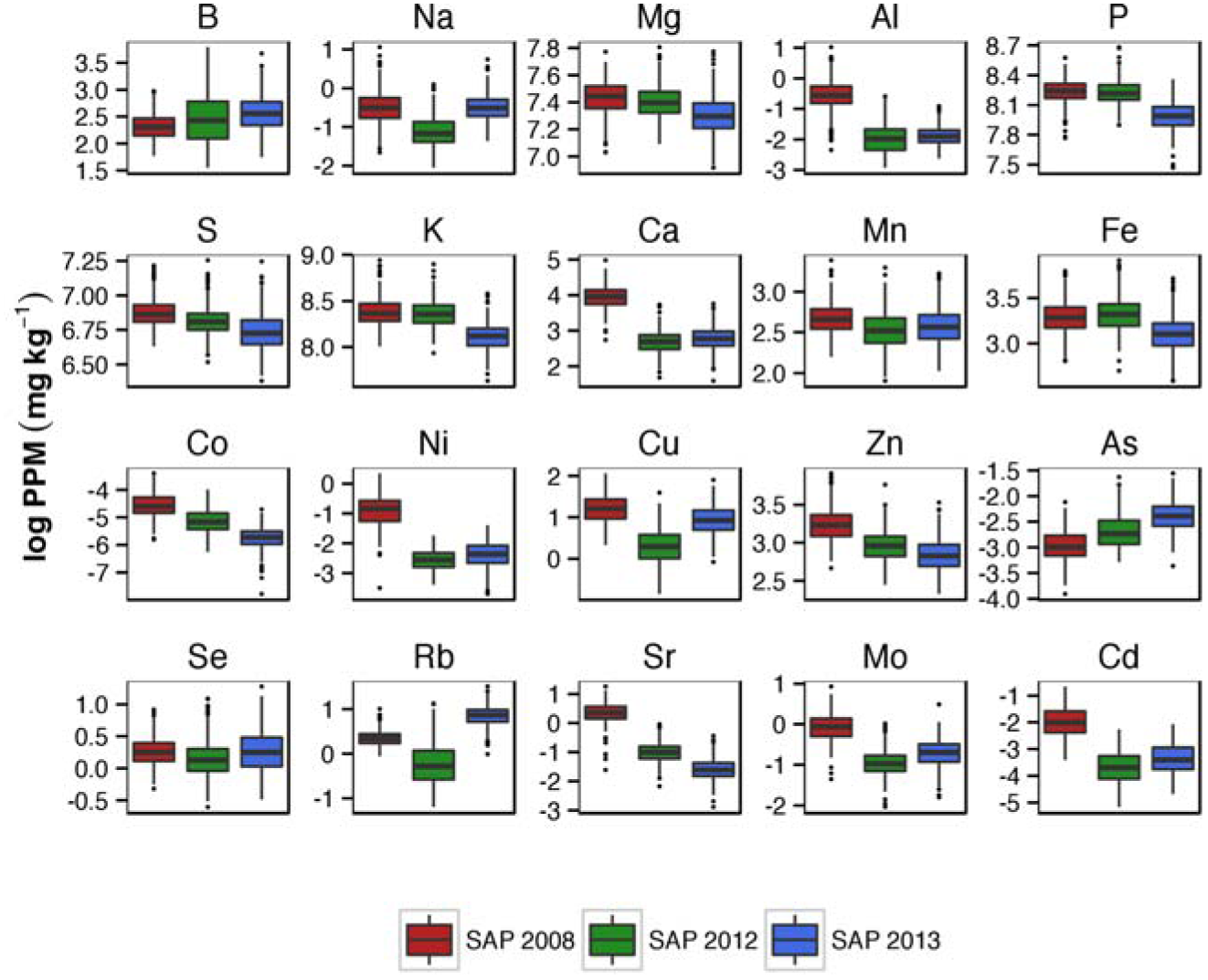
Box plots with median, minimum and maximum values, and interquartile ranges for the 20 elements in three SAP experimental populations. The raw concentration values for each of the elements were log transformed to obtain normally distributed phenotypes.

We used two different approaches to identify the shared regulation of elemental accumulation. Pairwise correlations were calculated and graphed (Supplemental table 4 and Figure 2a), and principal component analysis (PCA) was carried out (Figure 2b). Highly correlated element pairs in our data included Mg-P, Mg-Mn, P-S and Mg-S. Divalent cations Ca^2+^ and Sr^2+^ are chemical analogs and strong correlation was observed between these two elements, consistent with previous reports in other species (Queen et al., 1963; Hutchin and Vaughan, 1968; Ozgen et al., 2011; Broadley and White, 2012). In the SAP, the first two principal components accounted for a large fraction of the phenotypic covariance (36%). Clustering of elements reflected known elemental relationships, including the covariation of Ca and Sr (Figure 2). A cluster of the essential metal micronutrients, Fe, Zn and Cu is distinguishable suggesting that their accumulation can be affected by a shared mechanism. Similarly clustering of Mg and P is consistent with previous studies in wheat (Peleg et al., 2009). Seed P is predominately stored as the Mg2+ salt of phytic acid (inositol-hexaphosphate; IP6), which may explain the significant positive correlation of these elements (Maathuis, 2009; Marschner and Marschner, 2012).

**Figure 2A.**
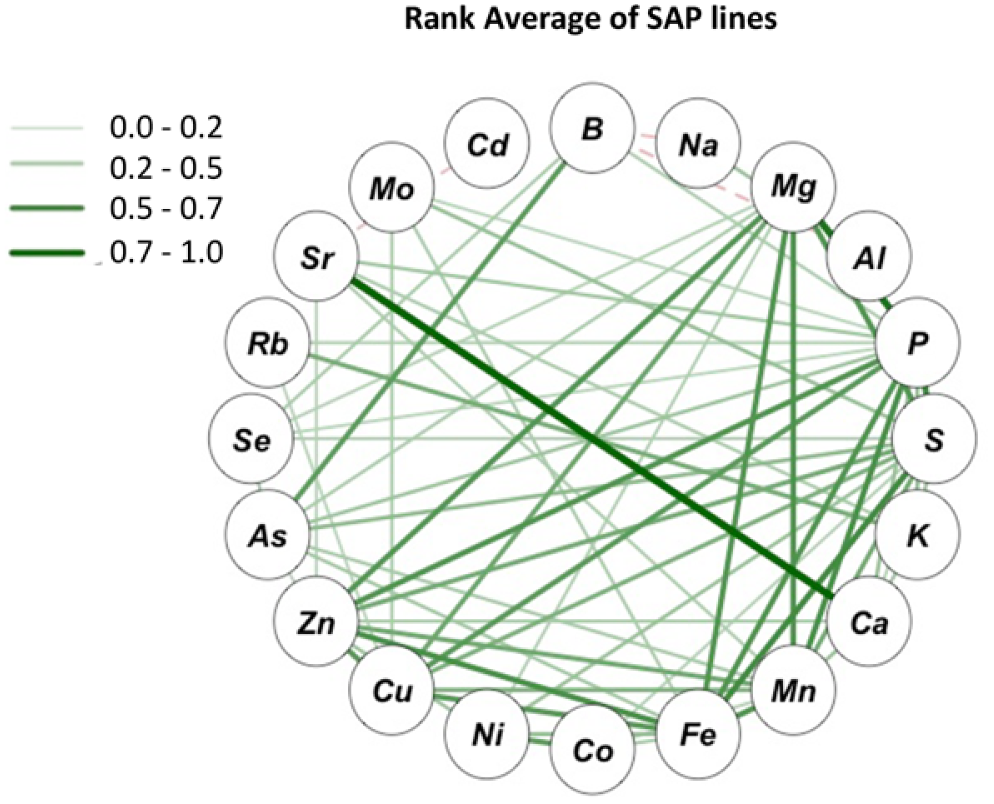
Correlation network of seed element concentrations using rank average data calculated across replicates from SAP association panels. Green solid lines represent positive correlation values. Red dashed lines represent negative correlation values. Intensity and thickness of lines indicate degree of correlation. Element correlation values can be found in Supplemental table 4. Correlation networks for SAP 2008, SAP 2012, and SAP 2013 can be found in Supplemental table 3.

**Figure 2B.**
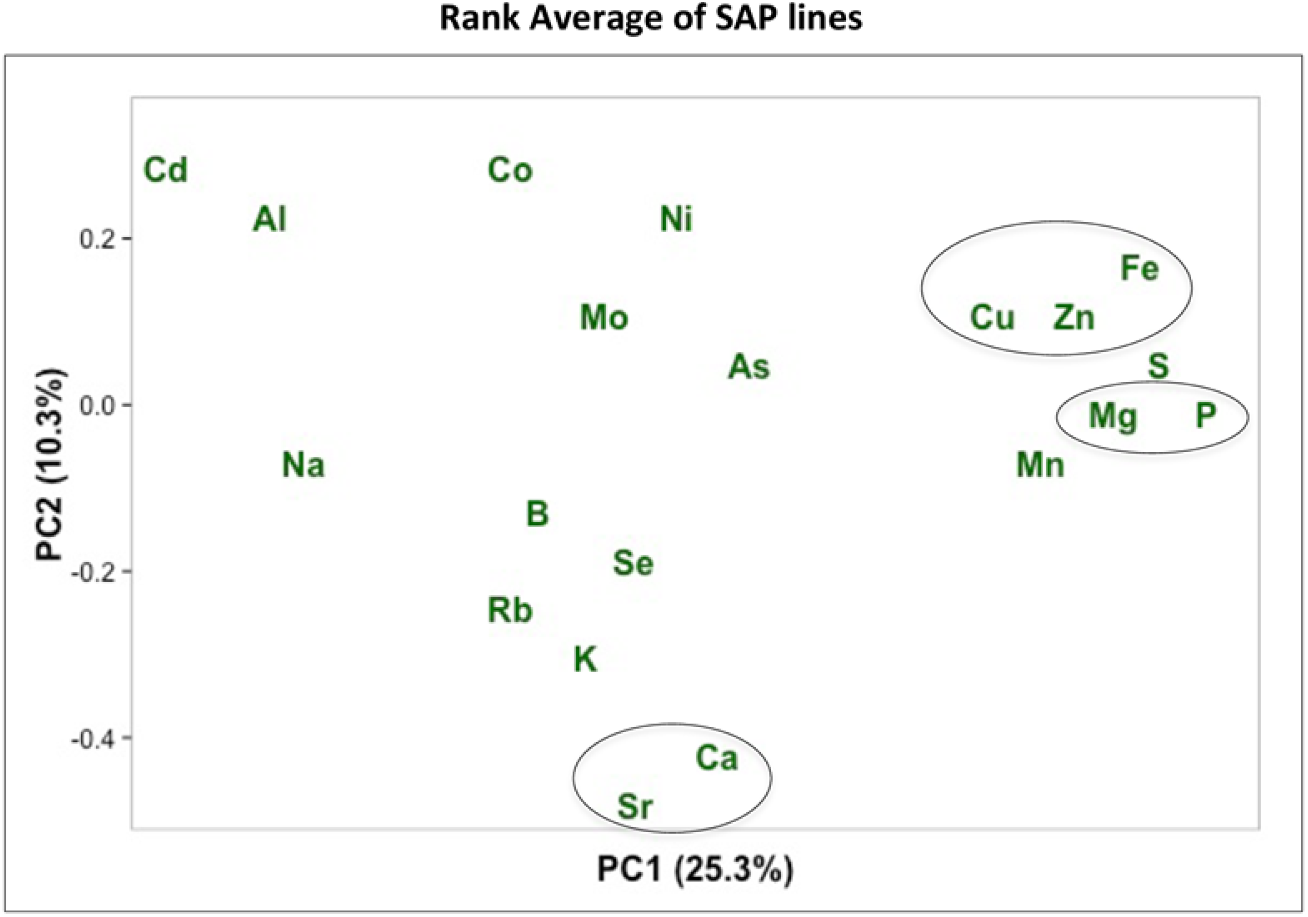
Principal component analysis applied to the rank average seed concentrations for 20 elements in the SAP lines across experiments. Each symbol represents a single element. PCA analysis for SAP 2008, SAP 2012 and SAP 2013 can be found in Supplemental table 4. Outlined elements reflect clustering of known elemental relationships

### Genome-wide association mapping of seed element traits

To dissect the genetic basis of natural variation for seed element concentration in sorghum seed, GWA mapping was performed using both an optimal model obtained from the multi-locus mixed model (MLMM) algorithm and a compressed mix linear model (CMLM) that accounts for population structure. For the MLMM analysis we considered several models to account for population structure as well as two different models to determine how many cofactors to add into the analysis (see Methods and Supplemental figure 5). We decided to use the kinship model to account for population structure and the most conservative mBonf model for selecting cofactors. We also used the conservative, Bonferroni-corrected threshold (P = 0.05) for CMLM, and identified overlapping SNPs significantly associated with seed element concentration using both approaches (Supplemental tables 5 and 6). Compared to traditional single-locus approaches (e.g. CMLM), MLMM utilizes multiple loci in the model, which contribute to a higher detection power and lower potential of false discoveries (Segura et al., 2012). MLMM also identified additional associations of interest. Significant SNPs identified with the MLMM approach were prioritized for further analysis (Supplemental table 5).

In an effort to comprehensively identify significant SNPs associated with element concentration, we created several datasets for GWA analysis. After averaging the two SAP 2013 growouts, each location was treated as an individual experiment. To link SAP experiments across environments, we ranked the individual lines of each experiment by element concentration and derived a robust statistic describing element accumulation for GWAS by using the average of ranks across the four SAP environments. By utilizing rank-order, we eliminated skewness and large variation in element concentration due to environmental differences (Conover and Iman, 1981). GWA scans across individual experiments identified 270, 228, and 207 significant SNPs for all twenty element traits in the SAP2008, SAP2012 and SAP2013 panels, respectively. In total we identified 255 significant loci in the ranked dataset for the twenty element traits (Supplemental table 5). The number of significant SNPs per element trait ranged from two (B) to 33 (Ca) and roughly reflected their heritabilities.

We identified several SNPs common to multiple environments (Supplemental table 8). For example, GWA for Ca concentration in all three of our SAP experiments identified significant SNPs within 5kb of locus Sobic.001G094200 on chromosome 1. Sobic.001G094200 is a putative calcium homeostasis regulator (CHoR1) (Zhang et al., 2012). We also identified several significant SNPs that colocalized for multiple element traits (Figure 3 and Supplemental table 9). Several of these SNPs were detected as significantly associated with multiple elements that are known to be coordinately regulated (Yi and Guerinot, 1996; Vert et al., 2002; Connolly et al., 2003; Lahner et al., 2003), and implicate candidate genes involved in regulation of multiple elements. For example, a SNP on chromosome 1 (S1_18898717) was a significant peak in both Mg and Mn GWA analysis (Figure 3). This SNP peak is in LD with the Arabidopsis homolog of AT3G15480. AT3G15480 is a protein of unknown function, however T-DNA knockout lines display mutant phenotypes in both Mg and Mn accumulation (www.ionomicshub.org, SALK_129213, Tray 449). T-DNA knockout lines in Arabidopsis also validated the significant peak for Co accumulation in the present study (S2_8464347). This SNP is linked to the homolog of AT5G63790, a NAC domain containing protein that imparts a significantly decreased Co phenotype in the T-DNA knockout line (www.ionomicshub.org,SALK_030702, Tray 1137).

**Figure 3.**
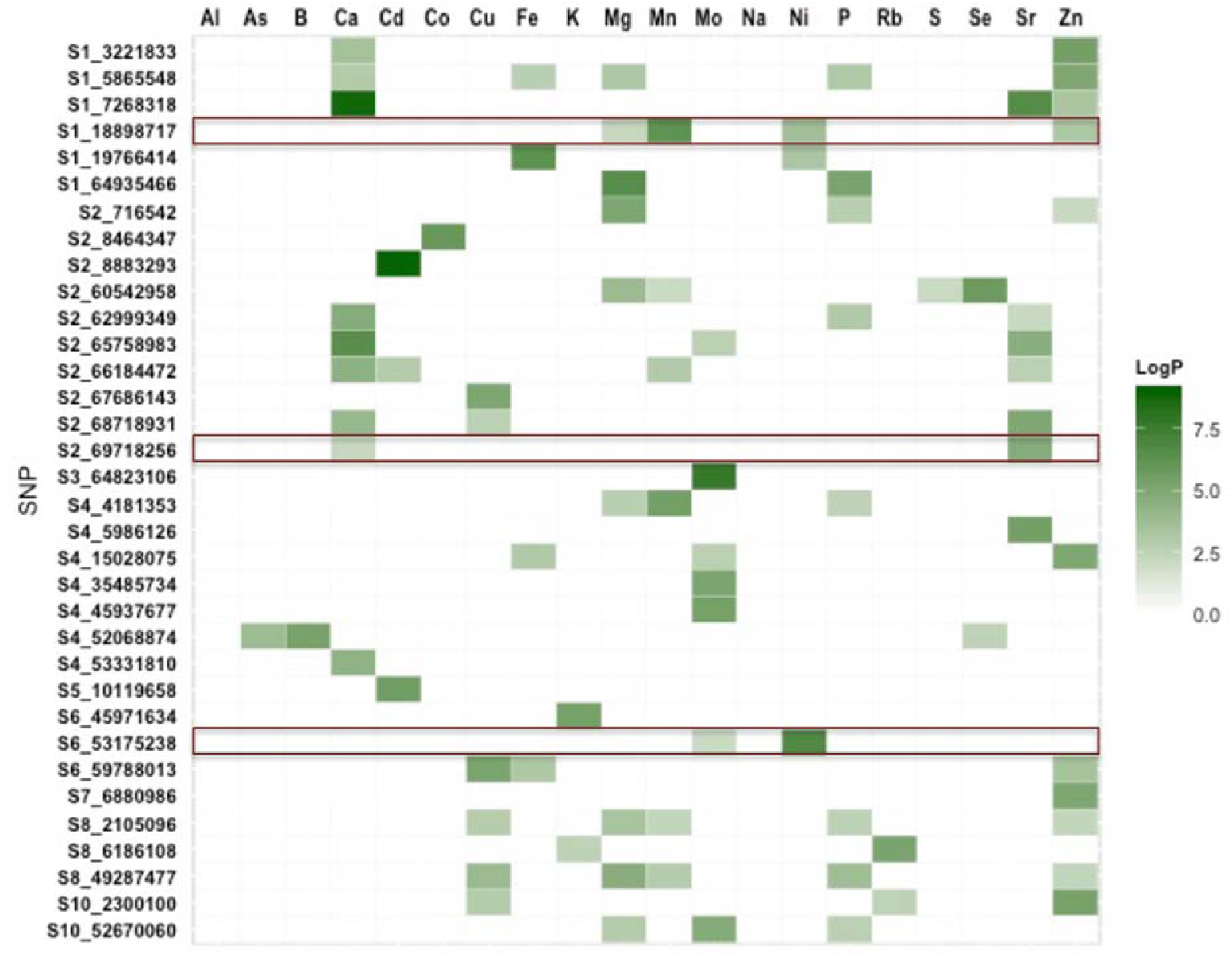
Heatmap displaying the Log P values of shared significant SNPs across 20 elements in the rank average dataset. Significance values below 2 are white and the ranges from 2.01 to 9.01 are shown in green (light to dark green color). Outlined in red are biologically relevant SN that colocalized for multiple elements.

We focused our interpretation efforts on the GWA results from the SAP ranked dataset, as these are the most likely to provide the tools to manipulate seed element content across multiple environments. The GWA results for each element trait obtained at the optimal step of the MLMM model were compiled. The data for Cd using the SAP ranked dataset is presented in Figure 4 as an example of the analysis procedure. GWA across multiple environments identified one significant SNP (S2_8883293) associated with Cd levels. (Figure 4A). The distribution of expected vs. observed P values, QQ-plots (Figure 4B and Supplemental figure 5), suggests that population structure was well controlled and false positive association signals were minimized using the kinship matrix plus cofactors.

**Figure 4A.**
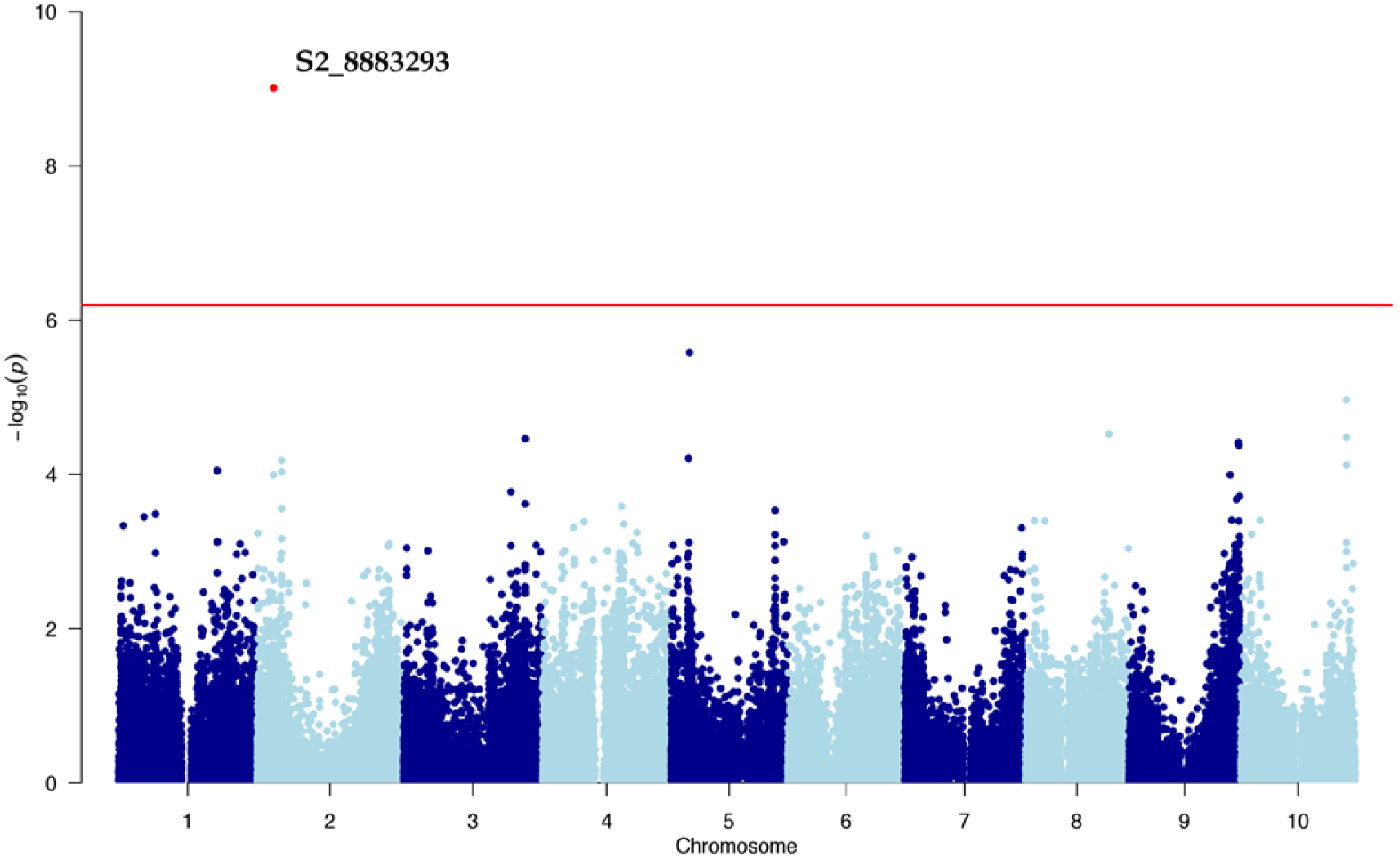
Manhattan plot displaying Cd GWAS results (—log_10_(P)) for the 10 sorghum chromosomes (x-axis) and associated *p* values for each marker (y-axis). The red lines indicate a Bonferroni-corrected threshold of 0.05.

**Figure 4B.**
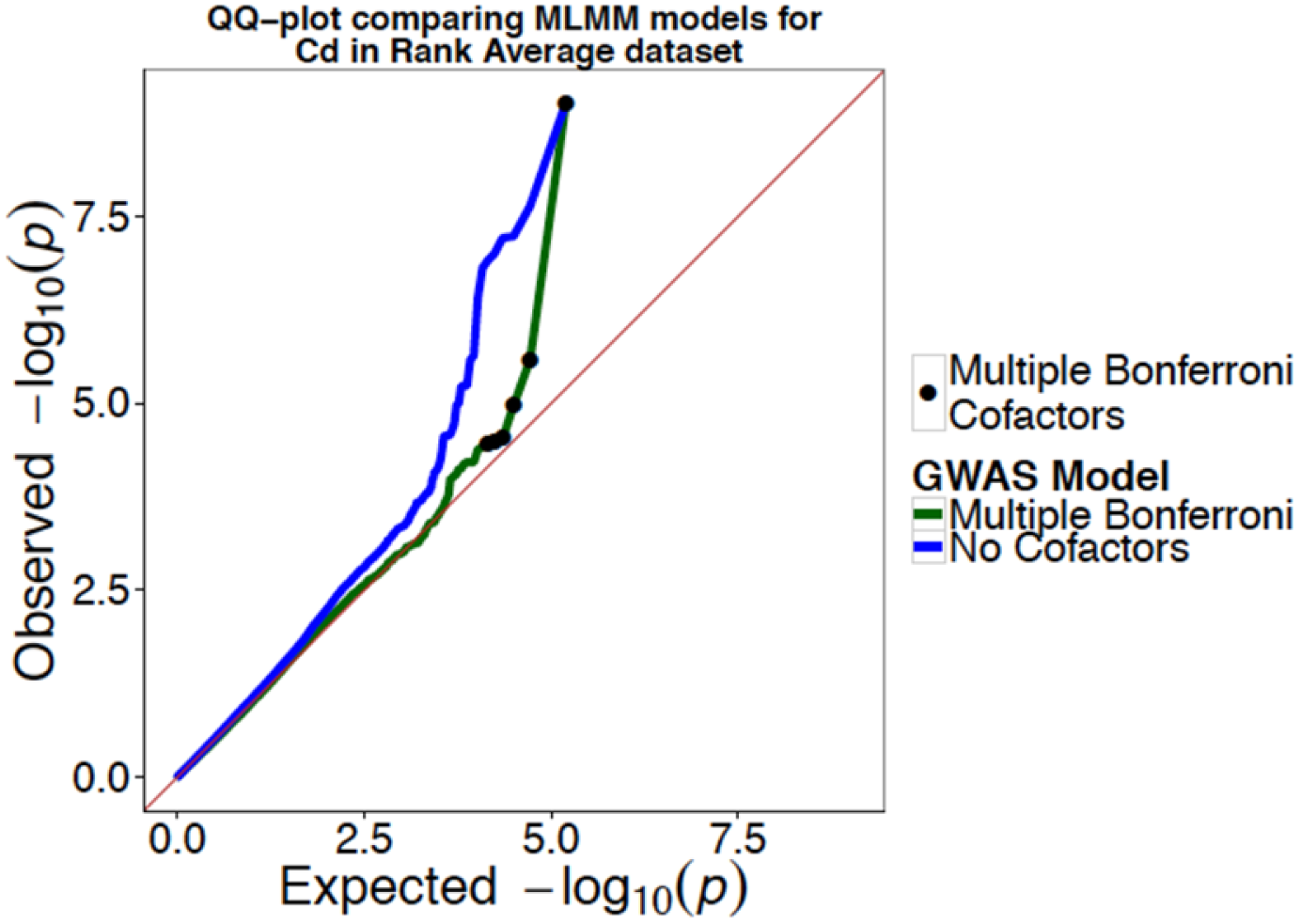
Quantile-quantile (QQ) of observed *p* values against the expected *p* values from the GWAS analysis for Cd element concentration. The MLMM mixed model includes cofactors that reduce inflation of *p* values (green line). The null model that does not consider significant cofactors, indicating the presence of *p* value inflation (blue line). The grey line indicates the expected *p* value distribution under the null hypothesis.

The optimal MLMM model (mBonf) included one SNP on chromosome 2, S2_8883293, that explained 18% of the phenotypic variation in cadmium (Figure 4C), and the allelic effects of each genotype were estimated (Figure 4D). The major-effect locus on chromosome 2 is in LD with a homolog of a well-characterized cadmium transporter in plants, heavy metal ATPase 2 (HMA2).

**Figure 4C.**
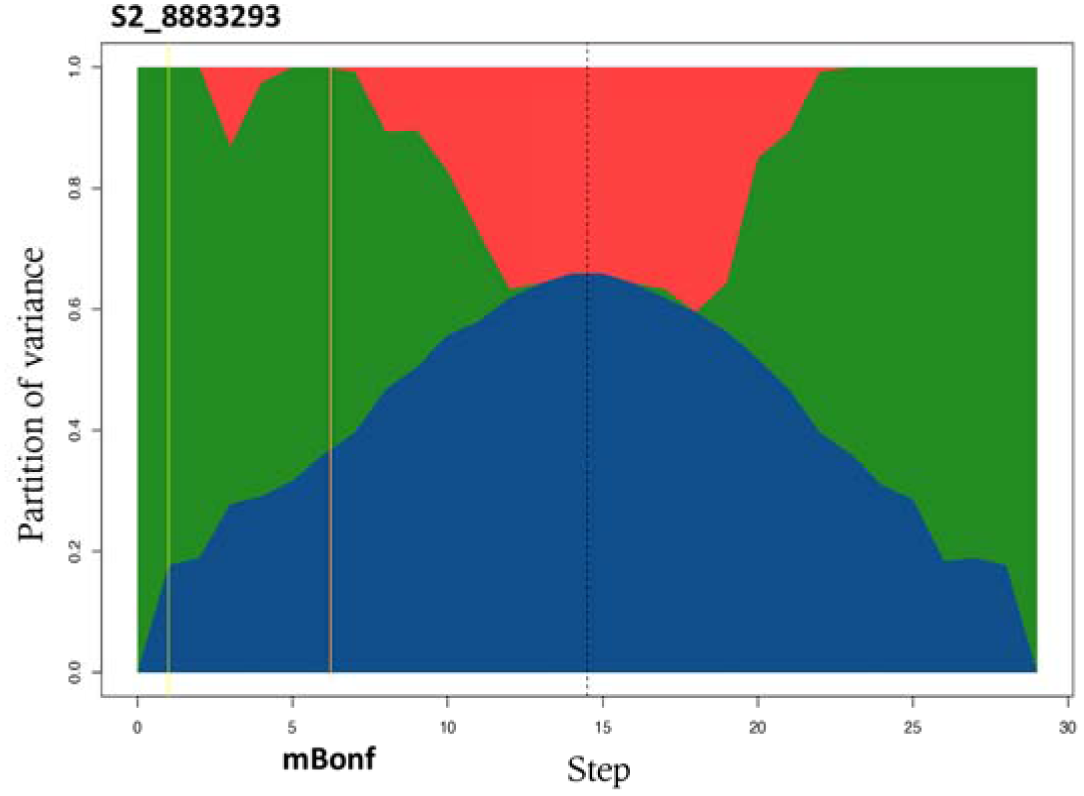
Evolution of genetic variance at each step of the MLMM (blue, genetic variance explained; green, total genetic variance; red, error). The yellow line indicates the variance with the inclusion of S2_8883293. The orange line indicates the optimal model selected by the multiple bonferroni criterion (mBonf).

**Figure 4D.**
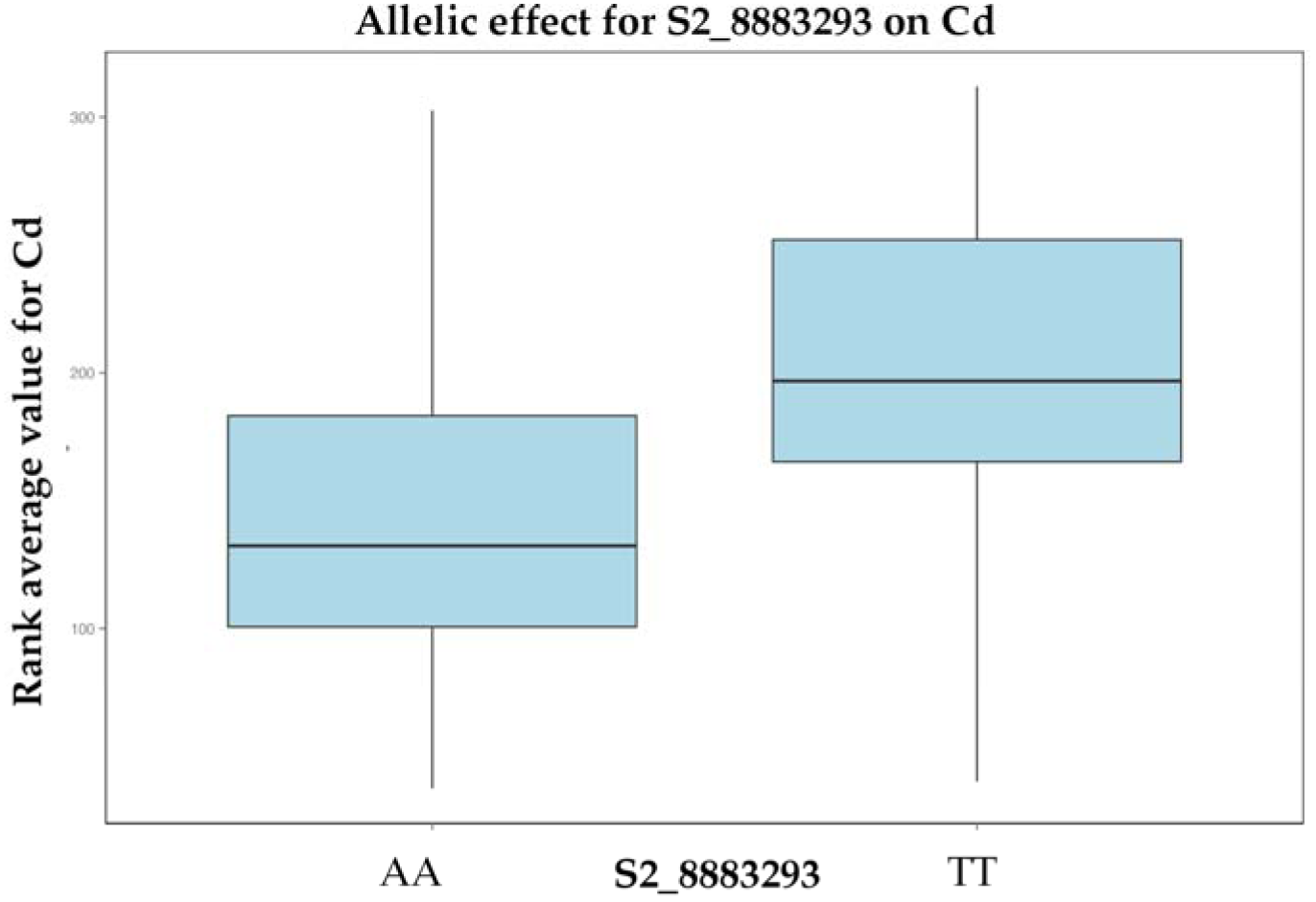
Allelic effect for the Cd significant SNP marker on chromosom 2.

## Discussion

### Ionome profiling for improved sorghum seed quality

Increasing the concentration of elements essential for human and animal nutrition (e.g. Fe and Zn) while simultaneously minimizing and increasing tolerance to anti-nutrients and toxic elements (e.g. As, Cd and Al) is a significant goal of fundamental research directed towards global crop improvement (Schroeder et al., 2013). Element homeostasis in plants, is affected by genotype, environment, soil properties, and nutrient interactions (Gregorio et al., 2000). While determining strategies to enhance or reduce element content for food or fuel, several components of seed element traits must be considered. These include: the heritability of the various element traits, genotype by environment interactions, and the availability of high-throughput element content screening tools (Ortiz-Monasterio et al., 2007). Differences in seed organic composition can also have large effects on the element composition of seeds, as different seed compartments will contain elements in different proportions. Variation in seed composition together with variation in sorghum seed sizes, violate the assumption of a uniform elemental concentration inherent in simple weight normalizations. Our data were not well modeled by a simple weight normalization (Supplemental figure 1), and we subsequently employed a rank transformation of the phenotypic data and linear model in the analysis (Ayana and Bekele, 2000; Baxter et al., 2014).

Our results demonstrate environmental effects on the range and means of element concentrations are largely element specific. In general, seed element concentrations did not exhibit large variation due to environmental effects. This contributed to high heritabilities for several elements and homeostasis of individual element concentration across very diverse environments (Figure 1 and Table 1). The high heritabilities for these traits demonstrate the feasibility of breeding strategies for the improvement of sorghum for seed element accumulation. Further, due to the known genetic contributors to trait covariation, selection strategies can include alteration of multiple traits, phenotypic correlations between traits or counter selection for undesirable traits (e.g. As accumulation). The high heritability and the relationships we report between important element elements, including Fe and Zn are encouraging for the development of breeding schema for improved element profiles for the alleviation of human malnutrition. Observed correlations of several elements indicate that changes in one or more elements can simultaneously affect the concentration of other elements present in the seed (Figure 2A). However, the individual effects of particular alleles can deviate from this pattern.

Trait correlations and covariation were used to uncover genetic associations for multiple elements. Even without more complicated analyses, we detected colocalized effects on several element traits (Supplemental table 4 and Supplemental table 8). For example, several significant SNPs colocalized for the strongly correlated element pairs of Ca and Sr (*r*= 0.79) as well as Mg and P (*r*= 0.71). Shared SNPs and colocalization of significant loci across multiple element traits suggest the possibility of tightly-linked genes or genes with pleiotropic effects and has been documented in recent GWA studies, including experiments in tomato (Sauvage et al., 2014) and rice (Zhao et al., 2011). In the present analysis, we applied a conservative threshold in our MLMM implementation and identified SNPs from the most complex model in which the P values of cofactors were below a defined threshold of 0.05. We implemented stringent parameters to eliminate false positives, but also risked the elimination of true positives. To identify additional candidate SNPs for further investigation, these stringent parameters can be relaxed to include association signals below the threshold.

### Candidate genes

One of the primary goals of this study was to utilize GWA analyses to identify candidate genes and novel loci implicated in the genetic regulation of sorghum seed element traits. We identified numerous significant SNPs for all twenty element traits that currently do not associate with known elemental accumulation genes. Although it is likely that a small fraction of these SNPs are false positives, many more may be novel associations with as-yet undiscovered causal genes and merit further investigation. We did, however, identify several significant SNPs that fall directly within a characterized candidate gene or are in close proximity, or LD, with putative candidates.

### Zinc

Zinc deficiency is a critical challenge for food crop production that results in decreased yields and nutritional quality. Zinc-enriched seeds result in better seedling vigor and higher stress tolerance on Zn-deficient soils (Cakmak, 2008). Here we identify a strong candidate for genetic improvement of zinc concentration in the in sorghum seed, Sobic.007G064900, an ortholog of Arabidopsis ZIP5, zinc transporter precursor (AT1G05300) (Table 2). AT1G05300 is a member of the ZIP family of metal transporter genes, and overexpression lines of this gene display increased Zn accumulation in Arabidopsis (www.ionomicshub.org,35SZip5_2 _Tray 700).

**Table II.**
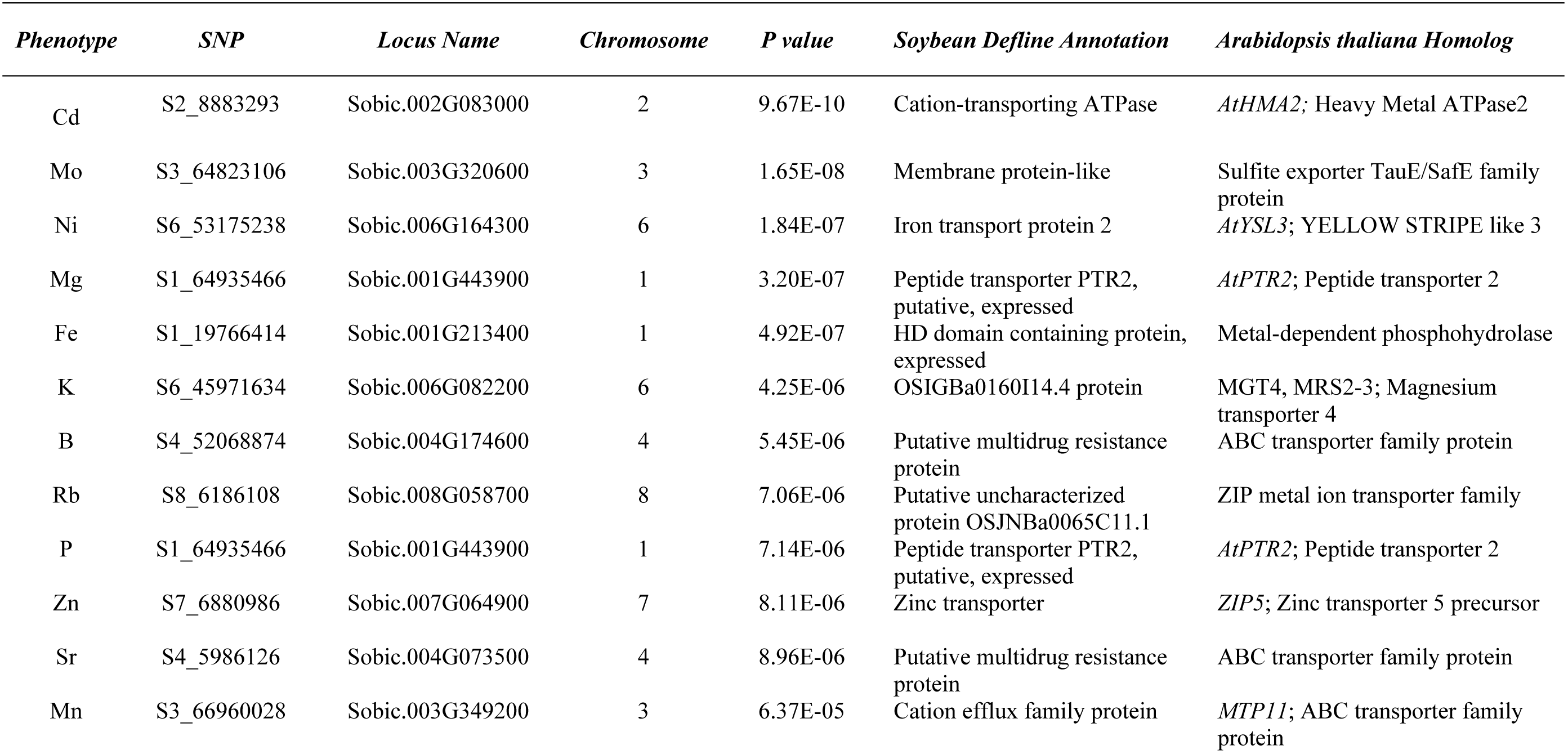
Detailed information for selected significant associations detected within the 20 element traits analyzed using the MLMM.

### Manganese

Associated with amino acid, lipid and carbohydrate metabolism, Mn is one of the essential elements critical to human and animal nutritional requirements (Aschner and Aschner, 2005). We identified significant GWAS associations in the putative sorghum homolog for member of the metal transporter encoding cation diffusion facilitator gene family MTP11 (Sobic.003G349200) (Table 2). The Arabidopsis ortholog, AtMTP11, confers Mn tolerance and transports Mn^2+^ via a proton-antiport mechanism in *Saccharomyces cerevisiae* (Delhaize et al., 2007).

### Cadmium

The seeds are a major source of essential nutrients, but can also be a source of toxic heavy metals, including cadmium. Contamination of ground water and subsequent uptake and absorption by the plant can result in high levels of Cd contamination in the seed (Arao and Ae, 2003). GWA analysis identified significant SNPs associated with a paralogous set of cation-transporting ATPases (Figure 4), Sobic.002G083000 and Sobic.002G083100. These are sorghum orthologs of Arabidopsis HMA genes in the heavy metal–transporting subfamily of the P-type ATPases. AtHMA3 participates in the vacuolar storage of Cd in Arabidopsis, and a recent study revealed that HMA3 is a major-effect locus controlling natural variation in leaf cadmium (Morel et al., 2009; Chao et al., 2012). The SNP alleles could be used immediately to potentially produce sorghum seed with lowered Cd2+ accumulation.

### Nickel

Ni is an essential nutrient required for plant growth. However, similar to Cd, high Ni concentrations in soil can be toxic to the plant, resulting in reduced biomass and crop yield. The most significant SNP for Ni concentration in the SAP 2008 environment (and present in SAP 2012 and the ranked dataset) was S6_53175238. This SNP is in LD with the candidate gene Sobic.006G164300, a homolog of the Yellow Stripe-Like 3 (YSL) family of proteins (Table 2). Originally identified in maize, the YSL proteins are a subfamily of oligopeptide transporters involved in metal uptake, homeostasis and long-distance transport (Curie et al., 2009). YSL3 is suggested to transport metals bound to nicotianamine (NA)(Curie et al., 2001) and in the metal hyperaccumulator *Thlaspi caerulescens* YSL3 functions as Ni–NA influx transporter (Gendre et al., 2007).

## Summary/Conclusion

In the present study, we utilized GWA mapping and rank transformation of the phenotypic data to scale GxE interactions and identify a number of genetic loci and candidate gene associations for immediate study and application to breeding strategies. The use of a multi-element, or ionomic approach, to the analysis allows for the identification of SNPs that confer multiple advantageous traits that can be selected for in breeding programs. We identify co-localization of significant SNPs for different elements, indicating potential coregulation through physiological processes of elemental uptake, transport, traffic and sequestration. Our results suggest that combining elemental profiling with GWA approaches can be useful for understanding the genetic loci underlying elemental accumulation and for improving nutritional content of sorghum. The data and analysis scripts used for this publication can be found at www.ionomicshub.org.

## Materials and Methods

### Plant material

The Sorghum Association Panel has been previously described (Casa et al., 2008). Seeds harvested from 407 lines that comprise the Sorghum Association Panel (SAP) were utilized for this study. The SAP 2008 seeds were obtained from Germplasm Resources Information Network (GRIN) and were produced in Lubbock, Texas by the USDA-ARS Cropping Systems Research Laboratory in 2008. The SAP 2012 seeds were produced in Puerto Vallarta, Mexico in 2012. The SAP 2013 seeds were produced in Florence, SC in 2013.

### Phenotypic Elemental Analysis

Four seeds per replicate were weighed from each individual and a minimum of two replicates from each line of the SAP 2008 and SAP 2013 panels were analyzed by ICP-MS. Each sample was digested with 2.5 mL of concentrated nitric acid at 95°C for 3 hours. Elemental analysis was performed with an ICP-MS for B, Na, Mg, Al, P, S, K, Ca, Mn, Fe, Co, Ni, Cu, Zn, As, Se, Rb, Sr, Mo and Cd following established protocols (Baxter et al., 2010). A reference sample derived from a pool of sorghum seed samples was generated and run after every 9^th^ sample to correct for ICP-MS run-to-run variation and within-run drift.

### Data Processing and Analysis

Phenotype data were generated for 407 SAP lines. GBS SNP markers for the SAP lines used in this study have been previously described (Morris et al., 2013). After removing SNPs with more than 20% missing data and minor allele frequencies below 0.05, genotype data for 78,012 SNPs remained. Broad sense heritability was calculated using the lmer function in the lme4 package to perform an analysis of variance with the experimental replicates of the SAP using previously described methods (Van Poecke et al., 2007; Bates et al., 2014). To ensure normality in the data distribution of the phenotype, the Box-Cox procedure was carried out on the phenotype scores to identify the best transformation method (Box and Cox, 1964). The ‘boxcox’ function in the MASS package in R was utilized to carry out the transformations (R Development Core Team, 2014; Ripley et al., 2014). In order to address potential confounding factors in the GWA analysis, specifically ICP run-to-run variation and the weight correction calculation, we used linear regression to compute residuals adjusted for ICP run and sample weight. These residuals were used to test for association with qualifying SNPs in the GWA analysis.

### GWAS

GWAS was executed in R using Genomic Association and Prediction Integrated Tool (GAPIT) using CMLM (Zhang et al., 2010; Lipka et al., 2012). Significant associations were determined by estimates of false discovery rate (FDR) (P = 0.05) (Benjamini and Hochberg, 1995). The CMLM uses a VanRaden kinship matrix and the first three principal components as covariates to account for population structure. MLMM is based on EMMA (Kang et al., 2008) and relies on the iterative use of a simple K, or Q+K, mixed-model algorithm. The kinship term, K, provides a fine-grained estimate of familial relatedness between lines. In addition, GWAS models often include a more granular measurement of population membership for each line, Q. To determine the necessity of using the more complex Q+K model to control for spurious allele associations, we analyzed QQ-plots generated from MLMM GWAS using a simple K model plus cofactors (Supplemental figure 5) and phenotypic distributions across known subpopulations (Supplemental figure 2). Phenotypic distributions across subpopulations were similar, indicating that population structure does not play a strong role in elemental accumulation. The QQ-plots indicate that after the addition of major effect loci to the model as cofactors, the *p* value distribution does not deviate drastically from the expected uniform distribution. These results indicate that for MLMM the mixed model containing only the kinship matrix, K, plus cofactors is sufficient to control for spurious allele associations due to population structure and cryptic relatedness.

At each step of the MLMM, the phenotypic variance is divided into genetic, random and explained variance. The most significant marker is included as a cofactor, and the variance components of the model are recalculated. With each successive iteration, the remaining genetic variance approaches zero, and an optimal model including cofactors that explains the genetic fraction of the phenotypic variance is determined. The MLMM method selects two models using stop criteria determined by two test statistics termed the multiple-Bonferroni criterion (mBonf) and the extended Bayesian information criterion (extBIC). The mBonf criteria selects a model wherein all cofactors have a *p* value below a Bonferroni-corrected threshold (Segura et al., 2012) and, in our experiments, this was the more stringent of the two model selection criteria (i.e. it favored less complex models) and was used for all further analyses. In addition, the genetic variance partition, described above, provides an estimate of heritability, termed pseudoheritability (Kang et al., 2010; Segura et al., 2012), explained by the model at each step. The missing heritability can be calculated from the model at the optimal step (mBonf). The percent variance explained by the model is the difference between the genetic variance at step 0 and the optimal step (Supplemental table 7).

The MLMM method utilized the multiple-Bonferroni criterion (mBonf) which selects a model wherein all cofactors have a *p* value below a Bonferroni-corrected threshold (Segura et al., 2012). We utilized a genome-wide significance threshold of *p* < 0.05 for the Bonferroni correction. A kinship matrix was constructed to correct for population structure and cryptic relatedness (Supplemental table 10). The kinship matrix was estimated from all of the SNPs in the dataset using the VanRaden method (VanRaden, 2008) in GAPIT (Lipka et al., 2012). Kinship was included as a random effect in the MLMM model.

## Supplemental Material

**Supplemental figure 1:** Correlation between seed weight and elemental concentration

**Supplemental figure 2:** Element distribution across sorghum subpopulations

**Supplemental figure 3:** Correlation network of seed element concentrations

**Supplemental figure 4:** PCA applied to the average seed concentrations for 20 elements

**Supplemental figure 5:** QQ Plots comparing MLMM models

**Supplemental table 1:** SAP lines

**Supplemental table 2:** Summary of one-way ANOVA for each evaluated trait and environment

**Supplemental table 3:** Means and standard deviations of seed element concentrations

**Supplemental table 4:** Correlation coefficients among seed element concentrations

**Supplemental table 5:** Significant SNPS identified by MLMM

**Supplemental table 6:** Significant SNPS identified by CMLM

**Supplemental table 7:** Calculations from MLMM-Pseudoheritability

**Supplemental table 8:** Shared significant SNPs across SAP datasets

**Supplemental table 9:** Significant SNPs shared for multiple elements

**Supplemental table 10:** Kinship matrix estimated from all of the SNPs

## Acknowledgments

This project was partially funded by the iHUB Visiting Scientist Program (http://www.ionomicshub.org), Chromatin, Inc., NSF EAGER (1450341) to I.B. and B.P.D., NSF IOS 1126950 to I.B., NSF IOS-0919739 to E.C., and BMGF (OPP 1052924) to B.P.D.

## Availability of supporting data

The datasets supporting the results of this article are available through Purdue Ionomics Information Management System (PiiMS) at http://www.ionomicshub.org

## Authors’ contributions

NS, GZ and IB wrote the manuscript, carried out ionomics assays, data analysis and interpretation of the results. ZB and RB contributed to experimental design and participated in tissue sampling. BD, EC, and SK participated in data analysis and interpretation of the results. All authors read, revised and approved the final manuscript.

## Parsed Citations

**Amaducci S, Monti A, Venturi G (2004) Non-structural carbohydrates and fibre components in sweet and fibre sorghum as affected by low and normal input techniques. Industrial Crops and Products 20: 111–118**

Pubmed: <u>Author and Title</u>

CrossRef: <u>Author and Ttle</u>

Google Scholar: <u>Author Only Ttle Only Author and Ttle</u>

**Arao T, Ae N (2003) Genotypic variations in cadmium levels of rice grain. Soil Science and Plant Nutrition 49: 473–479**

Pubmed: <u>Author and Ttle</u>

CrossRef: <u>Author and Ttle</u>

Google Scholar: <u>Author Only Ttle Only Author and Ttle</u>

**Aschner JL, Aschner M (2005) Nutritional aspects of manganese homeostasis. Molecular aspects of medicine 26: 353–362**

Pubmed: <u>Author and Ttle</u>

CrossRef: <u>Author and Ttle</u>

Google Scholar: <u>Author Only Ttle Only Author and Ttle</u>

**Ayana A, Bekele E (2000) Geographical patterns of morphological variation in sorghum (Sorghum bicolor (L.) Moench) germplasm from Ethiopia and Eritrea: quantitative characters. Euphytica 115: 91–104**

Pubmed: <u>Author and Ttle</u>

CrossRef: <u>Author and Ttle</u>

Google Scholar: <u>Author Only Ttle Only Author and Ttle</u>

**Bates D, Mächler M, Bolker B, Walker S (2014) Fitting linear mixed-effects models using lme4. arXiv preprint arXiv:1406.5823**

Pubmed: <u>Author and Ttle</u>

CrossRef: <u>Author and Ttle</u>

Google Scholar: <u>Author Only Ttle Only Author and Ttle</u>

**Baxter I, Brazelton JN, Yu D, Huang YS, Lahner B, Yakubova E, Li Y, Bergelson J, Borevitz JO, Nordborg M (2010) A, coastal cline in sodium accumulation in Arabidopsis thaliana is driven by natural variation of the sodium transporter AtHKT1; 1. PLoS genetics 6 e1001193**

Pubmed: <u>Author and Ttle</u>

CrossRef: <u>Author and Ttle</u>

Google Scholar: <u>Author Only Ttle Only Author and Ttle</u>

**Baxter IR, Gustin JL, Settles AM, Hoekenga OA (2013) Ionomic characterization of maize kernels in the intermated B73x Mo17 population. Crop Science 53: 208–220**

Pubmed: <u>Author and Ttle</u>

CrossRef: <u>Author and Ttle</u>

Google Scholar: <u>Author Only Ttle Only Author and Ttle</u>

**Baxter IR, Vitek O, Lahner B, Muthukumar B, Borghi M, Morrissey J, Guerinot ML, Salt DE (2008) The leaf ionome as a multivariable system to detect a plant’s physiological status. Proceedings of the National Academy of Sciences 105: 12081–12086**

Pubmed: <u>Author and Ttle</u>

CrossRef: <u>Author and Ttle</u>

Google Scholar: <u>Author Only Ttle Only Author and Ttle</u>

**Baxter IR, Ziegler G, Lahner B, Mickelbart MV, Foley R, Danku J, Armstrong P, Salt DE, Hoekenga OA (2014) Single-Kernel Ionomic Profiles Are Highly Heritable Indicators of Genetic and Environmental Influences on Elemental Accumulation in Maize Grain (Zea mays). PLoS One 9: e87628**

Pubmed: <u>Author and Ttle</u>

CrossRef: <u>Author and Ttle</u>

Google Scholar: <u>Author Only Ttle Only Author and Ttle</u>

**Benjamani Y, Hochberg Y (1995) Controlling the false discovery rate: a practical and powerful approach to multiple testing. Journal of the Royal Statistical Society. Series B (Methodological): 289–300**

Pubmed: <u>Author and Ttle</u>

CrossRef: <u>Author and Ttle</u>

Google Scholar: <u>Author Only Ttle Only Author and Ttle</u>

**Box GE, Cox DR (1964) An analysis of transformations. Journal of the Royal Statistical Society. Series B (Methodological): 211–252**

Pubmed: <u>Author and Ttle</u>

CrossRef: <u>Author and Ttle</u>

Google Scholar: <u>Author Only Ttle Only Author and Ttle</u>

**Broadley MR, White PJ (2012) Some elements are more equal than others: soil-to-plant transfer of radiocaesium and radiostrontium, revisited. Plant and soil 355: 23–27**

Pubmed: <u>Author and Ttle</u>

CrossRef: <u>Author and Ttle</u>

Google Scholar: <u>Author Only Ttle Only Author and Ttle</u>

**Buescher E, Achberger T, Amusan I, Giannini A, Ochsenfeld C, Rus A, Lahner B, Hoekenga O, Yakubova E, Harper JF (2010) Natural genetic variation in selected populations of Arabidopsis thaliana is associated with ionomic differences. PLoS One 5: e11081**

Pubmed: <u>Author and Ttle</u>

CrossRef: <u>Author and Ttle</u>

Google Scholar: <u>Author Only Ttle Only Author and Ttle</u>

**Cakmak I (2008) Enrichment of cereal grains with zinc: agronomic or genetic biofortification? Plant and Soil 302: 1–17**

Pubmed: <u>Author and Ttle</u>

CrossRef: <u>Author and Ttle</u>

Google Scholar: <u>Author Only Ttle Only Author and Ttle</u>

**Casa AM, Pressoir G, Brown PJ, Mitchell SE, Rooney WL, Tuinstra MR, Franks CD, Kresovich S (2008) Community resources and strategies for association mapping in sorghum. Crop science 48: 30–40**

Pubmed: <u>Author and Ttle</u>

CrossRef: <u>Author and Ttle</u>

Google Scholar: <u>Author Only Ttle Only Author and Ttle</u>

**Chao D-Y, Silva A, Baxter I, Huang YS, Nordborg M, Danku J, Lahner B, Yakubova E, Salt DE (2012) Genome-wide association studies identify heavy metal ATPase3 as the primary determinant of natural variation in leaf cadmium in Arabidopsis thaliana. PLoS genetics 8: e1002923**

Pubmed: <u>Author and Ttle</u>

CrossRef: <u>Author and Ttle</u>

Google Scholar: <u>Author Only Ttle Only Author and Ttle</u>

**Connolly EL, Campbell NH, Grotz N, Prichard CL, Guerinot ML (2003) Overexpression of the FRO2 ferric chelate reductase confers tolerance to growth on low iron and uncovers posttranscriptional control. Plant Physiology 133: 1102–1110**

Pubmed: <u>Author and Ttle</u>

CrossRef: <u>Author and Ttle</u>

Google Scholar: <u>Author Only Ttle Only Author and Ttle</u>

**Conover WJ, Iman RL (1981) Rank transformations as a bridge between parametric and nonparametric statistics. The American Statistician 35: 124–129**

Pubmed: <u>Author and Ttle</u>

CrossRef: <u>Author and Ttle</u>

Google Scholar: <u>Author Only Ttle Only Author and Ttle</u>

**Curie C, Cassin G, Couch D, Divol F, Higuchi K, Le Jean M, Misson J, Schikora A, Czernic P, Mari S (2009) Metal movement within the plant: contribution of nicotianamine and yellow stripe 1-like transporters. Annals of botany 103: 1–11**

Pubmed: <u>Author and Ttle</u>

CrossRef: <u>Author and Ttle</u>

Google Scholar: <u>Author Only Ttle Only Author and Ttle</u>

**Curie C, Panaviene Z, Loulergue C, Dellaporta SL, Briat J-F, Walker EL (2001) Maize yellow stripe1 encodes a membrane protein directly involved in Fe (III) uptake. Nature 409: 346–349**

Pubmed: <u>Author and Ttle</u>

CrossRef: <u>Author and Ttle</u>

Google Scholar: <u>Author Only Ttle Only Author and Ttle</u>

**Das P, Samantaray S, Rout G (1997) Studies on cadmium toxicity in plants: a review. Environmental pollution 98: 29–36**

Pubmed: <u>Author and Ttle</u>

CrossRef: <u>Author and Ttle</u>

Google Scholar: <u>Author Only Ttle Only Author and Ttle</u>

**Delhaize E, Gruber BD, Pittman JK, White RG, Leung H, Miao Y, Jiang L, Ryan PR, Richardson AE (2007) A, role for the AtMTP11 gene of Arabidopsis in manganese transport and tolerance. The Plant Journal 51: 198–210**

Pubmed: <u>Author and Ttle</u>

CrossRef: <u>Author and Ttle</u>

Google Scholar: <u>Author Only Ttle Only Author and Ttle</u>

**Gendre D, Czernic P, Conéjéro G, Pianelli K, Briat JF, Lebrun M, Mari S (2007) TcYSL3, a member of the YSL gene family from the hyper-accumulator Thlaspi caerulescens, encodes a nicotianamine-Ni/Fe transporter. The Plant Journal 49: 1–15**

Pubmed: <u>Author and Ttle</u>

CrossRef: <u>Author and Ttle</u>

Google Scholar: <u>Author Only Ttle Only Author and Ttle</u>

**Gomez-Becerra HF, Yazici A, Ozturk L, Budak H, Peleg Z, Morgounov A, Fahima T, Saranga Y, Cakmak I (2010) Genetic variation and environmental stability of grain mineral nutrient concentrations in T riticum dicoccoides under five environments. Euphytica 171: 39–52**

Pubmed: <u>Author and Ttle</u>

CrossRef: <u>Author and Ttle</u>

Google Scholar: <u>Author Only Ttle Only Author and Ttle</u>

**Graham R, Senadhira D, Beebe S, Iglesias C, Monasterio I (1999) Breeding for micronutrient density in edible portions of staple food crops: conventional approaches. Field Crops Research 60: 57–80**

Pubmed: <u>Author and Ttle</u>

CrossRef: <u>Author and Ttle</u>

Google Scholar: <u>Author Only Ttle Only Author and Ttle</u>

**Gregorio GB, Senadhira D, Htut H, Graham RD (2000) Breeding for trace mineral density in rice. Food & Nutrition Bulletin 21: 382386**

Pubmed: <u>Author and Ttle</u>

CrossRef: <u>Author and Ttle</u>

Google Scholar: <u>Author Only Ttle Only Author and Ttle</u>

**Huang X, Wei X, Sang T, Zhao Q, Feng Q, Zhao Y, Li C, Zhu C, Lu T, Zhang Z (2010) Genome-wide association studies of 14 agronomic traits in rice landraces. Nature genetics 42: 961–967**

Pubmed: <u>Author and Ttle</u>

CrossRef: <u>Author and Ttle</u>

Google Scholar: <u>Author Only Ttle Only Author and Ttle</u>

**Hutchin ME, Vaughan BE (1968) Relation between simultaneous Ca and Sr transport rates in isolated segments of vetch, barley, and pine roots. Plant physiology 43: 1913–1918**

Pubmed: <u>Author and Ttle</u>

CrossRef: <u>Author and Ttle</u>

Google Scholar: <u>Author Only Ttle Only Author and Ttle</u>

**Kang HM, Sul JH, Service SK, Zaitlen NA Kong S-y, Freimer NB, Sabatti C, Eskin E (2010) Variance component model to account for sample structure in genome-wide association studies. Nature genetics 42: 348–354**

Pubmed: <u>Author and Ttle</u>

CrossRef: <u>Author and Ttle</u>

Google Scholar: <u>Author Only Ttle Only Author and Ttle</u>

**Kang HM, Zaitlen NA Wade CM, Kirby A, Heckerman D, Daly MJ, Eskin E (2008) Efficient control of population structure in model organism association mapping. Genetics 178: 1709–1723**

Pubmed: <u>Author and Ttle</u>

CrossRef: <u>Author and Ttle</u>

Google Scholar: <u>Author Only Ttle Only Author and Ttle</u>

**Kimber CT, Dahlberg JA Kresovich S (2013) The gene pool of Sorghum bicolor and its improvement. In Genomics of the Saccharinae. Springer, pp 23–41**

Pubmed: <u>Author and Ttle</u>

CrossRef: <u>Author and Ttle</u>

Google Scholar: <u>Author Only Ttle Only Author and Ttle</u>

**Lahner B, Gong J, Mahmoudian M, Smith EL, Abid KB, Rogers EE, Guerinot ML, Harper JF, Ward JM, McIntyre L (2003) Genomic scale profiling of nutrient and trace elements in Arabidopsis thaliana. Nature biotechnology 21: 1215–1221**

Pubmed: <u>Author and Ttle</u>

CrossRef: <u>Author and Ttle</u>

Google Scholar: <u>Author Only Ttle Only Author and Ttle</u>

**Lestienne I, Icard-Vernière C, Mouquet C, Picq C, Trèche S (2005) Effects of soaking whole cereal and legume seeds on iron, zinc and phytate contents. Food Chemistry 89: 421–425**

Pubmed: <u>Author and Ttle</u>

CrossRef: <u>Author and Ttle</u>

Google Scholar: <u>Author Only Ttle Only Author and Ttle</u>

**Lipka AE, Tian F, Wang Q, Peiffer J, Li M, Bradbury PJ, Gore MA, Buckler ES, Zhang Z (2012) GAPIT: genome association and prediction integrated tool. Bioinformatics 28: 2397–2399**

Pubmed: <u>Author and Ttle</u>

CrossRef: <u>Author and Ttle</u>

Google Scholar: <u>Author Only Ttle Only Author and Ttle</u>

**Ma JF, Yamaji N, Mitani N, Xu X-Y, Su Y-H, McGrath SP, Zhao F-J (2008) Transporters of arsenite in rice and their role in arsenic accumulation in rice grain. Proceedings of the National Academy of Sciences 105: 9931–9935**

Pubmed: <u>Author and Ttle</u>

CrossRef: <u>Author and Ttle</u>

Google Scholar: <u>Author Only Ttle Only Author and Ttle</u>

**Maathuis FJ (2009) Physiological functions of mineral macronutrients. Current opinion in plant biology 12: 250–258**

Pubmed: <u>Author and Ttle</u>

CrossRef: <u>Author and Ttle</u>

Google Scholar: <u>Author Only Ttle Only Author and Ttle</u>

**Marschner H, Marschner P (2012) Marschner’s mineral nutrition of higher plants, Vol 89. Academic press**

Pubmed: <u>Author and Ttle</u>

CrossRef: <u>Author and Ttle</u>

Google Scholar: <u>Author Only Ttle Only Author and Ttle</u>

**Mengesha MH (1966) Chemical composition of teff (Eragrostis tef) compared with that of wheat, barley and grain sorghum. Economic Botany 20: 268–273**

Pubmed: <u>Author and Ttle</u>

CrossRef: <u>Author and Ttle</u>

Google Scholar: <u>Author Only Ttle Only Author and Ttle</u>

**Monti A, Di Virgilio N, Venturi G (2008) Mineral composition and ash content of six major energy crops. Biomass and Bioenergy 32: 216–223**

Pubmed: <u>Author and Ttle</u>

CrossRef: <u>Author and Ttle</u>

Google Scholar: <u>Author Only Ttle Only Author and Ttle</u>

**Morel M, Crouzet J, Gravot A, Auroy P, Leonhardt N, Vavasseur A, Richaud P (2009) AtHMA3, a P1B-ATPase allowing Cd/Zn/Co/Pb vacuolar storage in Arabidopsis. Plant Physiology 149: 894–904**

Pubmed: <u>Author and Ttle</u>

CrossRef: <u>Author and Ttle</u>

Google Scholar: <u>Author Only Ttle Only Author and Ttle</u>

**Morris GP, Ramu P, Deshpande SP, Hash CT, Shah T, Upadhyaya HD, Riera-Lizarazu O, Brown PJ, Acharya CB, Mitchell SE (2013) Population genomic and genome-wide association studies of agroclimatic traits in sorghum. Proceedings of the National Academy of Sciences 110: 453–458**

Pubmed: <u>Author and Ttle</u>

CrossRef: <u>Author and Ttle</u>

Google Scholar: <u>Author Only Ttle Only Author and Ttle</u>

**Neucere NJ, Sumrell G (1980) Chemical composition of different varieties of grain sorghum. Journal of agricultural and food chemistry 28: 19–21**

Pubmed: <u>Author and Ttle</u>

CrossRef: <u>Author and Ttle</u>

Google Scholar: <u>Author Only Ttle Only Author and Ttle</u>

**Norton GJ, Deacon CM, Xiong L, Huang S, Meharg AA, Price AH (2010) Genetic mapping of the rice ionome in leaves and grain: identification of QTLs for 17 elements including arsenic, cadmium, iron and selenium. Plant and soil 329: 139–153**

Pubmed: <u>Author and Ttle</u>

CrossRef: <u>Author and Ttle</u>

Google Scholar: <u>Author Only Ttle Only Author and Ttle</u>

**Obernberger I, Biedermann F, Widmann W, Riedl R (1997) Concentrations of inorganic elements in biomass fuels and recovery in the different ash fractions. Biomass and bioenergy 12: 211–224**

Pubmed: <u>Author and Ttle</u>

CrossRef: <u>Author and Ttle</u>

Google Scholar: <u>Author Only Ttle Only Author and Ttle</u>

**Organization WH (2002) The world health report 2002: reducing risks, promoting healthy life. World Health Organization**

Pubmed: <u>Author and Ttle</u>

CrossRef: <u>Author and Ttle</u>

Google Scholar: <u>Author Only Ttle Only Author and Ttle</u>

**Ortiz-Monasterio J, Palacios-Rojas N, Meng E, Pixley K, Trethowan R, Pena R (2007) Enhancing the mineral and vitamin content of wheat and maize through plant breeding. Journal of Cereal Science 46: 293–307**

Pubmed: <u>Author and Ttle</u>

CrossRef: <u>Author and Ttle</u>

Google Scholar: <u>Author Only Ttle Only Author and Ttle</u>

**Ozgen S, Busse JS, Palta JP (2011) Influence of Root Zone Calcium on Shoot Tip Necrosis and Apical Dominance of Potato Shoot: Simulation of This Disorder by Ethylene Glycol Tetra Acetic Acid and Prevention by Strontium. HortScience 46: 1358–1362**

Pubmed: <u>Author and Ttle</u>

CrossRef: <u>Author and Ttle</u>

Google Scholar: <u>Author Only Ttle Only Author and Ttle</u>

**Peleg Z, Cakmak I, Ozturk L, Yazici A, Jun Y, Budak H, Korol AB, Fahima T, Saranga Y (2009) Quantitative trait loci conferring grain mineral nutrient concentrations in durum wheat× wild emmer wheat RIL population. Theoretical and Applied Genetics 119: 353–369**

Pubmed: <u>Author and Ttle</u>

CrossRef: <u>Author and Ttle</u>

Google Scholar: <u>Author Only Ttle Only Author and Ttle</u>

**Queen WH, Fleming HW, O’Kelley JC (1963) Effects on Zea mays seedlings of a strontium replacement for calcium in nutrient media. Plant physiology 38: 410**

Pubmed: <u>Author and Ttle</u>

CrossRef: <u>Author and Ttle</u>

Google Scholar: <u>Author Only Ttle Only Author and Ttle</u>

**R Development Core Team (2014) R: a language and environment for statistical computing. Vienna, Austria: R Foundation for Statistical Computing; 2012. Open access available at: http://cran.r-project.org**

Pubmed: <u>Author and Ttle</u>

CrossRef: <u>Author and Ttle</u>

Google Scholar: <u>Author Only Ttle Only Author and Ttle</u>

**Ragaee S, Abdel-Aal E-SM, Noaman M (2006) Antioxidant activity and nutrient composition of selected cereals for food use. Food Chemistry 98: 32–38**

Pubmed: <u>Author and Ttle</u>

CrossRef: <u>Author and Ttle</u>

Google Scholar: <u>Author Only Ttle Only Author and Ttle</u>

**Ripley B, Venables B, Bates DM, Hornik K, Gebhardt A, Firth D, Ripley MB (2014) Package ‘MASS’.**

**Salt DE, Baxter I, Lahner B (2008) Ionomics and the study of the plant ionome. Annu. Rev. Plant Biol. 59: 709–733**

Pubmed: <u>Author and Ttle</u>

CrossRef: <u>Author and Ttle</u>

Google Scholar: <u>Author Only Ttle Only Author and Ttle</u>

**Sauvage C, Segura V, Bauchet G, Stevens R, Do PT, Nikoloski Z, Fernie AR, Causse M (2014) Genome Wide Association in tomato reveals 44 candidate loci for fruit metabolic traits. Plant Physiology: pp. 114.241521**

Pubmed: <u>Author and Ttle</u>

CrossRef: <u>Author and Ttle</u>

Google Scholar: <u>Author Only Ttle Only Author and Ttle</u>

**Schroeder JI, Delhaize E, Frommer WB, Guerinot ML, Harrison MJ, Herrera-Estrella L, Horie T, Kochian LV, Munns R, Nishizawa NK (2013) Using membrane transporters to improve crops for sustainable food production. Nature 497: 60–66**

Pubmed: Author and Ttle

CrossRef: Author and Ttle

**Segura V, Vilhjálmsson BJ, Platt A, Korte A, Seren Ü, Long Q, Nordborg M (2012) An efficient multi-locus mixed-model approach for genome-wide association studies in structured populations. Nature genetics 44: 825–830**

Pubmed: <u>Author and Ttle</u>

CrossRef: <u>Author and Ttle</u>

Google Scholar: <u>Author Only Ttle Only Author and Ttle</u>

**Shi R, Li H, Tong Y, Jing R, Zhang F, Zou C (2008) Identification of quantitative trait locus of zinc and phosphorus density in wheat (Triticum aestivum L.) grain. Plant and soil 306: 95–104**

Pubmed: <u>Author and Ttle</u>

CrossRef: <u>Author and Ttle</u>

Google Scholar: <u>Author Only Ttle Only Author and Ttle</u>

**Šimic D, Drinic SM, Zdunic Z, Jambrovic A, Ledencan T, Brkic J, Brkic A, Brkic I (2012) Quantitative trait loci for biofortification traits in maize grain. Journal of Heredity 103: 47–54**

Pubmed: <u>Author and Ttle</u>

CrossRef: <u>Author and Ttle</u>

Google Scholar: <u>Author Only Ttle Only Author and Ttle</u>

**Tian F, Bradbury PJ, Brown PJ, Hung H, Sun Q, Flint-Garcia S, Rocheford TR, McMullen MD, Holland JB, Buckler ES (2011) Genome-wide association study of leaf architecture in the maize nested association mapping population. Nature genetics 43: 159–162**

Pubmed: <u>Author and Ttle</u>

CrossRef: <u>Author and Ttle</u>

Google Scholar: <u>Author Only Ttle Only Author and Ttle</u>

**USDA A, National Genetic Resources Program. Germplasm Resources Information Network - (GRIN) In, National Germplasm Resources Laboratory, Beltsville, Maryland.**

**Van Poecke RM, Sato M, Lenarz-Wyatt L, Weisberg S, Katagiri F (2007) Natural variation in RPS2-mediated resistance among Arabidopsis accessions: correlation between gene expression profiles and phenotypic responses. The Plant Cell Online 19: 4046–4060**

Pubmed: <u>Author and Ttle</u>

CrossRef: <u>Author and Ttle</u>

Google Scholar: <u>Author Only Ttle Only Author and Ttle</u>

**VanRaden P (2008) Efficient methods to compute genomic predictions. Journal of dairy science 91: 4414–4423**

Pubmed: <u>Author and Ttle</u>

CrossRef: <u>Author and Ttle</u>

Google Scholar: <u>Author Only Ttle Only Author and Ttle</u>

**Vert G, Grotz N, Dédaldéchamp F, Gaymard F, Guerinot ML, Briat J-F, Curie C (2002) IRT1, an Arabidopsis transporter essential for iron uptake from the soil and for plant growth. The Plant Cell Online 14: 1223–1233**

Pubmed: <u>Author and Ttle</u>

CrossRef: <u>Author and Ttle</u>

Google Scholar: <u>Author Only Ttle Only Author and Ttle</u>

**Vreugdenhil D, Aarts M, Koornneef M, Nelissen H, Ernst W (2004) Natural variation and QTL analysis for cationic mineral content in seeds of Arabidopsis thaliana. Plant, Cell & Environment 27: 828–839**

Pubmed: <u>Author and Ttle</u>

CrossRef: <u>Author and Ttle</u>

Google Scholar: <u>Author Only Ttle Only Author and Ttle</u>

**Waters BM, Grusak MA (2008) Quantitative trait locus mapping for seed mineral concentrations in two Arabidopsis thaliana recombinant inbred populations. New Phytologist 179: 1033–1047**

Pubmed: <u>Author and Ttle</u>

CrossRef: <u>Author and Ttle</u>

Google Scholar: <u>Author Only Ttle Only Author and Ttle</u>

**White PJ, Broadley MR (2005) Biofortifying crops with essential mineral elements. Trends in plant science 10: 586–593**

Pubmed: <u>Author and Ttle</u>

CrossRef: <u>Author and Ttle</u>

Google Scholar: <u>Author Only Ttle Only Author and Ttle</u>

**Yi Y, Guerinot ML (1996) Genetic evidence that induction of root Fe (III) chelate reductase activity is necessary for iron uptake under iron deficiency‡. The Plant Journal 10: 835–844**

Pubmed: <u>Author and Ttle</u>

CrossRef: <u>Author and Ttle</u>

Google Scholar: <u>Author Only Ttle Only Author and Ttle</u>

**Zhang M, Pinson SR, Tarpley L, Huang X-Y, Lahner B, Yakubova E, Baxter I, Guerinot ML, Salt DE (2014) Mapping and validation of quantitative trait loci associated with concentrations of 16 elements in unmilled rice grain. Theoretical and Applied Genetics 127: 137–165**

Pubmed: <u>Author and Ttle</u>

CrossRef: <u>Author and Ttle</u>

Google Scholar: <u>Author Only Ttle Only Author and Ttle</u>

**Zhang T, Zhao X, Wang W, Pan Y, Huang L, Liu X, Zong Y, Zhu L, Yang D, Fu B (2012) Comparative transcriptome profiling of chilling stress responsiveness in two contrasting rice genotypes. PloS one 7: e43274**

Pubmed: <u>Author and Ttle</u>

CrossRef: <u>Author and Ttle</u>

Google Scholar: <u>Author Only Ttle Only Author and Ttle</u>

**Zhang Z, Ersoz E, Lai C-Q, Todhunter RJ, Tiwari HK, Gore MA, Bradbury PJ, Yu J, Arnett DK, Ordovas JM (2010) Mixed linear model approach adapted for genome-wide association studies. Nature genetics 42: 355–360**

Pubmed: <u>Author and Ttle</u>

CrossRef: <u>Author and Ttle</u>

Google Scholar: <u>Author Only Ttle Only Author and Ttle</u>

**Zhao K, Tung C-W, Eizenga GC, Wright MH, Ali ML, Price AH, Norton GJ, Islam MR, Reynolds A, Mezey J (2011) Genome-wide association mapping reveals a rich genetic architecture of complex traits in Oryza sativa. Nature communications 2: 467**

Pubmed: <u>Author and Ttle</u>

CrossRef: <u>Author and Ttle</u>

Google Scholar: <u>Author Only Ttle Only Author and Ttle</u>

